# Drivers of mosquito presence and abundance in urban and garden ponds in a European city

**DOI:** 10.64898/2026.04.07.716928

**Authors:** Irene Tornero, Barbara Barta, Andrew J. Hamer, Zoltán Soltész, Thu-Hương Huỳnh, Ádám Mészáros, Zsófia Horváth

## Abstract

Mosquitoes in urban environments are considered unpleasant by citizens, and they can represent serious health risks as disease vectors. Invasive *Aedes* species are continuously spreading in Europe and have been increasingly reported from urban settlements. Some of these mosquito species have been associated with small artificial habitats. Therefore, their presence may also be expected in small plastic garden ponds, which are numerous in urban settings across many parts of Europe. Here, we aimed to determine whether urban waters host a high occurrence of mosquitoes and to identify the potential abiotic, biotic and landscape drivers of mosquito presence and abundance in urban ponds. We sampled 53 urban ponds in the city of Budapest (Hungary) of both natural and anthropogenic origin, and 40 garden ponds, which cover both highly urbanised areas and those with high green index in the suburbs. We collected data on macroinvertebrate communities, pond management, physico-chemical parameters, and pond characteristics (mainly morphology and vegetation cover) measured *in situ*. Two different mosquito detection techniques were used: dip-net sampling and eDNA. While only one invasive species (*Aedes koreicus*) was detected, occurring in a single pond, several other species were present, including potential malaria vectors increasingly reported from urban environments: the *Anopheles maculipennis* complex, *An. claviger* and *An. plumbeus*. Fish presence was negatively associated with both the presence and abundance of mosquitoes, regardless of pond type. Contrary to expectations, urbanisation did not play a major role in explaining mosquito presence or abundance. These results highlight the importance of local pond characteristics, and particularly the role of fish presence, in regulating mosquito populations in ponds in the urban landscape, although the broader ecological effects of fish on pond communities should also be considered.

## INTRODUCTION

Urbanisation leads to changes in biodiversity mainly through land use and land cover changes, which reduce both species richness and evenness for most biotic communities (Grimm et al., 2008). Green and blue spaces in urban areas (e.g., parks, forests, rivers and ponds) can buffer the negative impact of urbanisation on biodiversity (Hyseni et al., 2021). Blue infrastructure (i.e., rivers, streams, canals and ponds) provides essential services and benefits for the urban population (Haase et al., 2014) as well as supporting freshwater biodiversity. Ponds are abundant in the urban landscape but are rarely of natural origin (Horváth et al., 2025). These ecosystems often have poor water quality due to pollution and may support exotic species (Oertli & Parris, 2019). Among other negative consequences for biodiversity, urbanisation both fragments the landscape and creates barriers to organisms’ dispersal, which is key for the assemblage of aquatic communities (Hamer & McDonnell, 2008; Sanderson et al., 2005).

Urban ponds are small freshwater habitats in cities, encompassing several types, such as natural or semi-natural, ornamental, industrial, stormwater retention, reservoirs, temple and fish ponds. They are commonly located on public property or owned by the state, municipality, institutions or private companies (Horváth et al., 2025). Urban ponds can be up to a few hectares in size and made of concrete, stone, or natural substrate, with a variable shoreline that is usually simpler and less vegetated than natural ponds (Oertli & Parris, 2019). Garden ponds are a specific subset of urban ponds located within domestic gardens owned and managed by a private individual (Hamer et al., 2024). They commonly have an ornamental function (Hill et al., 2023; Oertli et al., 2023), are stocked with fish and have a small size (usually up to a few m^2^) (Hamer et al., 2024; Hassall, 2014; Hill et al., 2021). Although these characteristics may reduce their contribution to sustaining urban biodiversity, they may, however, contribute to local species richness by acting as stepping stones for the movement of insects and amphibians across the urban landscape (Gledhill et al., 2008; Hill et al., 2021). Due to their private nature, we have little information on how they are managed and their local diversity. Thus, there is still a need to further investigate their role in contributing to biodiversity, ecosystem functions, services and disservices (Horváth et al., 2025).

Mosquitoes are commonly associated with pond ecosystems, including those situated within urban landscapes (Knight et al., 2003). The general public often associates urban wetlands with health risks linked to mosquitoes (Hanford et al., 2019). Urbanisation is frequently linked to high mosquito abundance and diversity (Perrin et al., 2023a), suggesting that high anthropogenic pressures can promote mosquito occurrence (Boerlijst et al., 2023). Urbanisation can provide numerous man-made breeding sites and refugia for mosquitoes, including flooded underground pipes and drainage systems, which some species exploit even under suboptimal conditions, as well as providing a stable water source during the dry season through irrigation and leakage (Norris, 2004; Valdelfener, et al., 2018; Vora, 2008). However, urbanisation may also negatively affect mosquito dispersal by posing barriers (Perrin et al., 2023b). In fact, a recent meta-analysis by Perrin et al. (2022) revealed an overall decline of mosquito presence in response to urbanisation, possibly linked to less availability of aquatic habitats (e.g., lower number of tree holes, ditches, vernal pools and leaf axils; Ferraguti et al., 2016; Gardner et al., 2014; Loaiza et al., 2019; Steiger et al., 2016). Furthermore, several studies have shown that urban ponds tend to host fewer mosquitoes than natural or rural ponds (see Ferraguti et al., 2016; Ibañez-Justicia et al., 2015; Versteirt et al., 2013). Therefore, the association between urbanisation and mosquito incidence is still not fully understood, given the inconsistent evidence in the literature.

Mosquitoes are organisms of major public health concern (World Health Organization, 2002). Several species of mosquitoes are of particular interest because they act as vectors for severe diseases. Several *Anopheles* species (Diptera, Culicidae) exhibit variable susceptibility and capability to transmit *Plasmodium* spp., and they occur widely throughout Europe. However, their detailed distribution patterns remain incompletely known (Tagliapietra et al., 2019). The most widely distributed belong to the *Anopheles maculipennis* complex, with several species of variable susceptibility to infection by *Plasmodium* spp. Other species, such as *An. algeriensis*, *An. claviger*, *An., hyrcanus*, *An. plumbeus* and *An. superpictus* have historically played a minor role as secondary vectors, although their vectorial competence is being reevaluated (Piperaki & Daikos, 2016). In particular, *An. plumbeus* was found to be susceptible to *Plasmodium falciparum*, thus raising suspicion of its involvement in cryptic malaria transmission in Central and Western Europe (Schaffner et al., 2012). Moreover, *Aedes koreicus*, another potential vector of mosquito-borne diseases, is also present in Europe (Brugueras et al., 2020). This species is already established in several European countries, including Hungary (European Centre for Disease Prevention and Control and European Food Safety Authority, 2023). Although *Ae. koreicus* has been associated with the outbreaks of both Japanese encephalitis and *Dirofilaria*, its role as a vector for these diseases remains unconfirmed (Schaffner et al., 2013). Therefore, reliably detecting mosquitoes in urban aquatic habitats is increasingly important. A variety of techniques are used to detect mosquitoes, including both active and passive approaches such as collecting mosquitoes in their natural habitats, deploying traps for adult mosquitoes, citizen science approaches to detect and locate them, and, more recently, using environmental DNA (eDNA) from water samples (Wittwer et al., 2024). The reliability and efficiency of the eDNA-based approach to detect mosquitoes, including the invasive species, have been shown in several studies (Schneider et al., 2016; Sakata et al., 2022; Gutiérrez-López et al., 2023).

Several abiotic and biotic factors can modulate mosquito populations in ponds. For instance, water quality parameters, such as sulphate concentration, have been associated with increased mosquito presence and abundance (Gadzama et al., 2018). Also, some studies found that mosquito abundance depends on habitat size, although both positive and negative relationships have been reported (Juliano, 2009; Sunahara et al., 2002). Due to their proximity to human activity, urban ponds tend to accumulate pollutants from surface runoff (Hassall, 2014; Oertli & Parris, 2019) and often have low water quality (Power et al., 2018). The presence of such water pollutants has been linked to biodiversity loss (Villalobos-Jiménez et al., 2016). In particular, aquatic invertebrates are sensitive to heavy metals, as demonstrated by both field and laboratory studies (Malaj et al., 2012). In addition to the unintentional input of contaminants, management practices in garden ponds, including the regular use of pesticides and/or algaecides (Hamer et al., 2024; Loram et al., 2011; Márton et al., 2025), may also negatively affect the biota. Among the taxa that could be potentially affected, several macroinvertebrate taxa are known predators of mosquito larvae (e.g., Culler & Lamp, 2009; Knight et al., 2003; Banerjee et al., 2010). Some amphibians also prey on mosquito larvae (Rubbo et al., 2011) and sometimes compete with them for the use of similar resources (Blaustein & Margalit, 1996). Moreover, the presence of fish, often inhabiting in garden ponds, negatively influences the abundance of several macroinvertebrate taxa (Thornhill, 2013). Thus, fish can exert a double biotic control, since they prey on mosquito larvae and on other natural predators of mosquito larvae (Becker et al., 2010; Horváth et al., 2025).

Given the frequent assumption that urban waters provide breeding habitats for mosquitoes, here, we aim to assess whether these secondary habitats in the urban landscape indeed host a large number of mosquitoes, including invasive or potential vectors of public health concern, in a Central European city. We also aimed to identify the most important environmental and landscape-level predictors of mosquito presence and abundance in these urban ponds, and compare results based on dip-net and eDNA samples. To this end, we sampled a large number of ponds across the Budapest metropolitan area (Hungary).

## METHODS

### Pond selection

We selected garden ponds from the “MyPond” project online survey (see Márton et al., 2025). First, all ponds were selected within a 36 km (which represents around 1.5 times the city’s radius and covers about twice the area of Budapest) radius circle from the centre of Budapest, and the level of urbanisation was calculated (see below). Then, ponds were categorised based on the level of urbanisation and binned into 10 main categories. From each category, a random set of 6 ponds was selected, and their owners were contacted. Two garden ponds (GW23 and GW24) that we sampled were agricultural ponds. Thus, they were not so representative of the pond type, but we kept them in the analysis to provide more complete data on urban mosquito occurrences. This resulted in 40 garden ponds in total that we could sample. For the rest of urban ponds, we aimed to sample all the ponds located within the Budapest metropolitan area, resulting in 53 urban ponds, which represent a close to extensive set of all such ponds in the city. Therefore, our dataset was constituted by a total of 93 ponds (Figure 1).

**Fig. 1.**
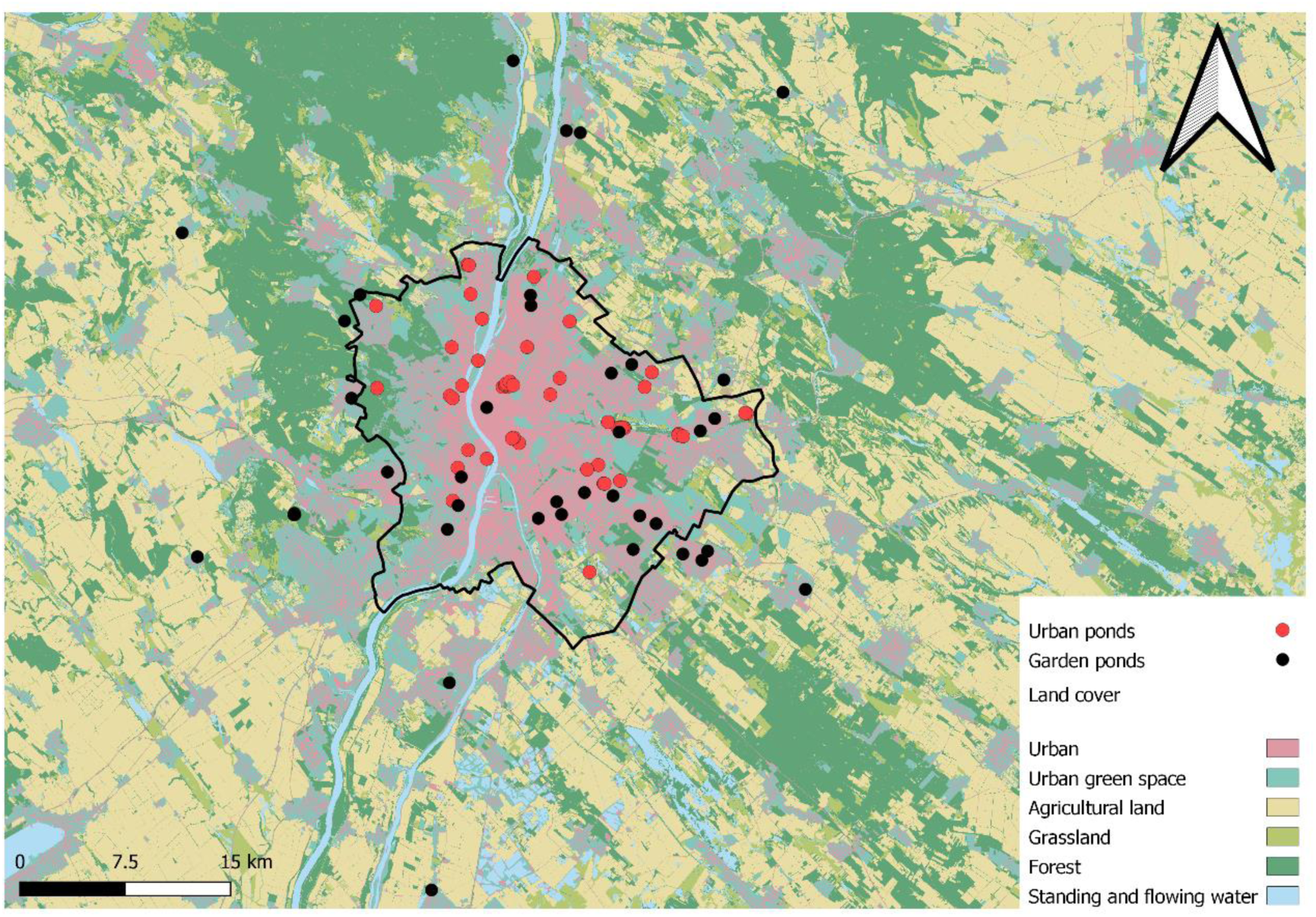
Map of the distribution of the 53 urban (red) and 40 garden (black) ponds that were sampled across the Budapest metropolitan area (Hungary). Land cover types as defined by ‘The Ecosystem Map of Hungary’ are shown.

### Sampling

Ponds were sampled between 4^th^ April and 17^th^ May 2022. Aquatic macroinvertebrate samples were collected by performing dip-net sweeps, where sampling effort (number of sweeps) was adjusted to pond size (Table S1 and S2). Sweeps were equally distributed across the different microhabitats present in each pond. Sampling was done from the shoreline in order to avoid damaging and altering the aesthetic appearance of these ornamental ponds. Samples were collected using an aquarium net (15 x 12 cm, mesh size: 500 µm) in the case of garden ponds, and a rectangular-framed (25 x 20 cm, mesh size: 500 µm) hand net in the case of urban ponds. Macroinvertebrate samples were preserved in 70% alcohol. In the laboratory, individuals were sorted, counted and identified to species level whenever possible, or to the lowest taxonomic category. The main identification keys used were Tachet et al. (2003), Nilsson (1996) and Becker et al. (2010). Later on, individual counts were standardised since samples were collected using nets of different sizes.

In parallel, eDNA from water samples was collected in the same ponds. Water was collected from ten different points in the ponds using a sterile ladle. Once homogenised, water was filtered using Sterivex filters (0.45 µm) and a sterile syringe. The samples were dried, preserved in 99% ethanol and kept in cold (-20 °C) until processing. DNA was extracted with the DNeasy Blood & Tissue Kits (QIAGEN), following the manufacturer’s protocol, with an additional ZYMO PCR Inhibitor removal step. Library preparation and sequencing were conducted by EnviroDNA (Australia). Mosquito DNA was amplified using a Culicidae-specific COI metabarcoding assay (Krol et al., 2019). Prior to full analysis, several candidate assays (COI, 16S, and 28S) were evaluated for specificity and taxonomic resolution, and the COI assay was selected as it provided the best performance for Culicidae detection in a pilot assay. Libraries were prepared using a two-step PCR protocol and sequenced on Illumina platforms (iSeq 100 and NextSeq 2000). Sequence data were quality-filtered and clustered into amplicon sequence variants using VSEARCH (Rognes et al., 2016). Taxonomic assignment was performed against a curated reference database derived from NCBI using a 95% similarity threshold. Negative extraction and PCR controls were included throughout library preparation to monitor contamination.

### Pond features

At each pond, surface area (m^2^), open water (visual estimate of the % of open water not covered by emergent or floating vegetation) and pond age were recorded (based on information from pond owners or managers). In the case of urban ponds, their area was either calculated using the Google Maps measuring tool or measured *in situ*. For garden ponds, we approximated the pond surface area based on the lengths and widths (that pond owners provided in the online survey; see Márton et al., 2025), by multiplying these measurements. Regarding pond age, in the case of garden ponds, if owners did not provide a specific year of installation but stated that they inherited the pond from a previous owner and provided the year they moved in, then pond age was calculated from that year (Hamer et al., 2024). For urban ponds, we used the ‘Historical Imagery’ from Google Earth Pro to establish when they appeared. Conductivity, pH and chlorophyll-a were measured *in situ* with portable meters (HANNA (HI 9142N), WTW (3620 IDS) and AquaPen AP 110-C, Photon System Instruments). Concentrations of TP (µg/L), TN (mg/L), TOC (mg/L), and heavy metals (Al, As, Ba, Cr, Cu, Fe, Li, Mn, Ni and Zn, in µg/L) as potential pollutants in urban settings were measured from water samples following the protocols from Eaton et al. (2005). The type of pond (garden or urban) was considered as a factor. This set of variables was considered as local abiotic predictors in the subsequent analyses.

The presence of fish and amphibians in each pond was also recorded. These two binary variables (1, presence; 0, absence) were considered as local biotic predictors of mosquito presence and abundance in the subsequent analyses. We estimated the abundance of those macroinvertebrate taxa that can potentially prey on mosquito larvae (for instance, Dytiscidae, Chaoboridae, Heteropterans, Odonates, etc.) and considered it as a local biotic variable (hereafter, abundance of predatory macroinvertebrates).

### Landscape-level data

To characterise the level of urbanisation surrounding the ponds, we used the Ecosystem Map of Hungary (project KEHOP-430-VEKOP-15-2016-00001, Ministry of Agriculture, 2019). The number of raster pixels (20×20 m) was counted within a 1 km radius circle around each focal pond using the Zonal Histogram tool. The classification of land cover was used as defined by The Ecosystem Map of Hungary and then these were grouped together into larger categories of ‘Forest’, ‘Grassland’, ‘Agricultural land’, ‘Standing and flowing waters’ and ‘Urban’ land cover types. The urban category contained pixels of tall and short buildings, sealed and dirt roads, railways and other artificial surfaces. Urbanisation was calculated as the proportion of total pixels within the 1 km radius covered by the ‘Urban’ land cover type (Oertli & Parris, 2019). All spatial analysis was carried out in QGIS 3.28.1 (QGIS Development Team, 2022).

For the analysis of spatial community structure, we used Moran’s Eigenvector Maps (MEMs) based on the spatial position of the ponds, using the “adespatial” R package (Dray et al., 2025). We only kept the positive MEM eigenvectors for the subsequent analyses.

### Statistical analysis

To reduce dimensionality and summarise the variation in heavy metal concentrations, we performed a Principal Component Analysis (PCA) on log-transformed data. The first principal component explaining the highest variance (40.4%) was retained for further analysis and used as an explanatory variable in subsequent analysis (Figure S1 in the Supplementary material). Hereafter, we will refer to this PC1 as ‘metalsPC1’.

Stepwise selection of positive MEMs was performed using a redundancy analysis (RDA) framework on the detrended mosquito community data set (Borcard et al., 2018). The selection process, based on the Akaike Information Criterion (AIC) and using both forward and backward stepwise search, identified the positive MEMs most strongly related to mosquito community composition (i.e., MEM2). We conducted a variation partitioning analysis to divide the variation of mosquito community composition among three sets of predictors: local abiotic, local biotic and landscape. Mosquito community data were Hellinger transformed. Local abiotic matrix was composed of pond features that were selected with a stepwise model selection with the ‘ordistep’ function from the complete set of local abiotic variables (i.e., open water, metalsPC1, TN and pond area). The local biotic matrix was composed of both amphibians and fish presence, and the abundance of predatory macroinvertebrates. Finally, urbanisation and the MEM2 constituted the landscape matrix. Variation partitioning was carried out using the ‘varpart’ function, while the significance of unique and shared fractions was tested using the function ‘anova’ in the R package vegan (Oksanen et al., 2025).

To analyse which predictors had the strongest effect on mosquito abundance, a linear model (LM) was used. Prior to this, a stepwise model selection was performed to select significant local abiotic variables to be included in the LM (“stepAIC” function, direction= “both”). The stepwise model selection identified pond area and metalsPC1 as the local abiotic variables best explaining mosquito abundance. Using the same procedure, the positive MEMs with a significant effect on mosquito abundance were selected (i.e., MEM2 and MEM5) and then included in the LM. The set of local biotic variables (amphibians and fish presence, and abundance of predatory macroinvertebrates) was also included. Similarly, to assess which predictors most strongly influenced mosquito presence based on dip-net samples, we fitted a generalised linear model (GLM) with a binomial distribution and logit link function. Prior to this, local abiotic variables and positive MEMs, separately, were selected using the ‘stepAIC’ function, which resulted in keeping metalsPC1 and open water, and MEM2, respectively. An analogous GLM was also fitted to the mosquito presence data derived from eDNA samples in order to compare the results obtained with the other sampling technique. Again, a prior selection of the positive MEMs and of the local abiotic variables was done, which resulted in keeping MEM17 and open water, respectively, for the GLM.

The three regression models were repeated, including the interaction term between fish presence and abundance of predatory macroinvertebrates, to test the possible effect of the biotic interaction between the mosquito predators.

## RESULTS

From the 93 ponds sampled, a total of 21 ponds hosted mosquitoes, of which 9 were garden ponds, and 12 were urban ponds based on dip-net sampling (see Table 1). Based on eDNA sampling, 8 garden and 6 urban ponds had mosquitoes. In 11 cases, eDNA detected Culicidae in samples where dip-net samples did not. Conversely, in 18 ponds, Culicidae were collected with dip-net sampling, but not detected by eDNA. Dip-net samples resulted in a higher total number of species detected, while the eDNA-based species were a subset of these (Figure 2). Dip-net samples detected 1 taxon shared between the garden and urban ponds, with 8 taxa unique to garden ponds and 7 taxa unique to urban ponds (Figure 2a). eDNA samples detected 2 taxa shared between the garden and urban ponds, with 2 taxa unique to garden ponds and only 1 taxon unique to urban ponds (Figure 2b). Both techniques detected mosquitoes in the same ponds on only three occasions (Figure 3a and Figure 3b). This suggests that the two sampling methods are complementary. We identified 14 species from five genera. In ponds where mosquitoes were present, their species numbers ranged between 1 and 4. Both detection techniques detected the same taxa on 3 occasions (Figure 3c). Thirteen taxa were exclusively detected by dip-netting whereas only two taxa were exclusively detected by eDNA method (Figure 3c). Among the three most invasive *Aedes* species in Europe, we only found *Aedes koreicus*, occurring in two urban ponds (Figure 4). Regarding potential malaria-vectors, *Anopheles maculipennis* was detected in five other public ponds and one garden pond, while two other lower-risk *Anopheles* species, *An. claviger* and *An. plumbeus*, were each found in three ponds. Of the five genera that we detected, eDNA samples detected all of them except *Ochlerotatus* (Figure 5).

**Fig. 2.**
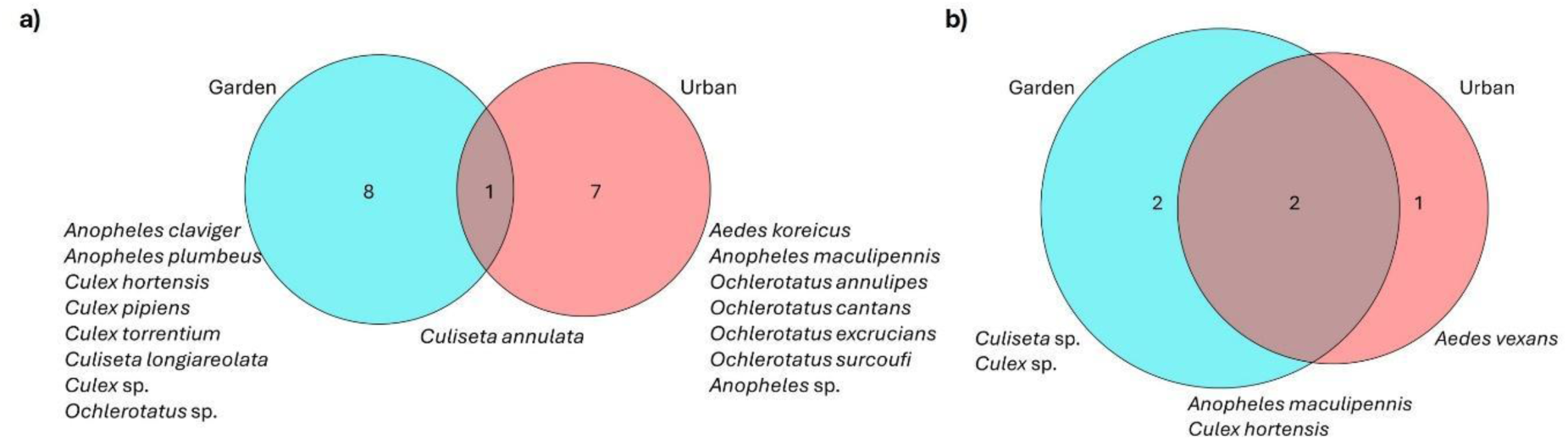
Venn diagrams showing the overlap in mosquito taxa detected among pond types. a) Data based on dip-netting; b) data based on eDNA.

**Fig. 3.**
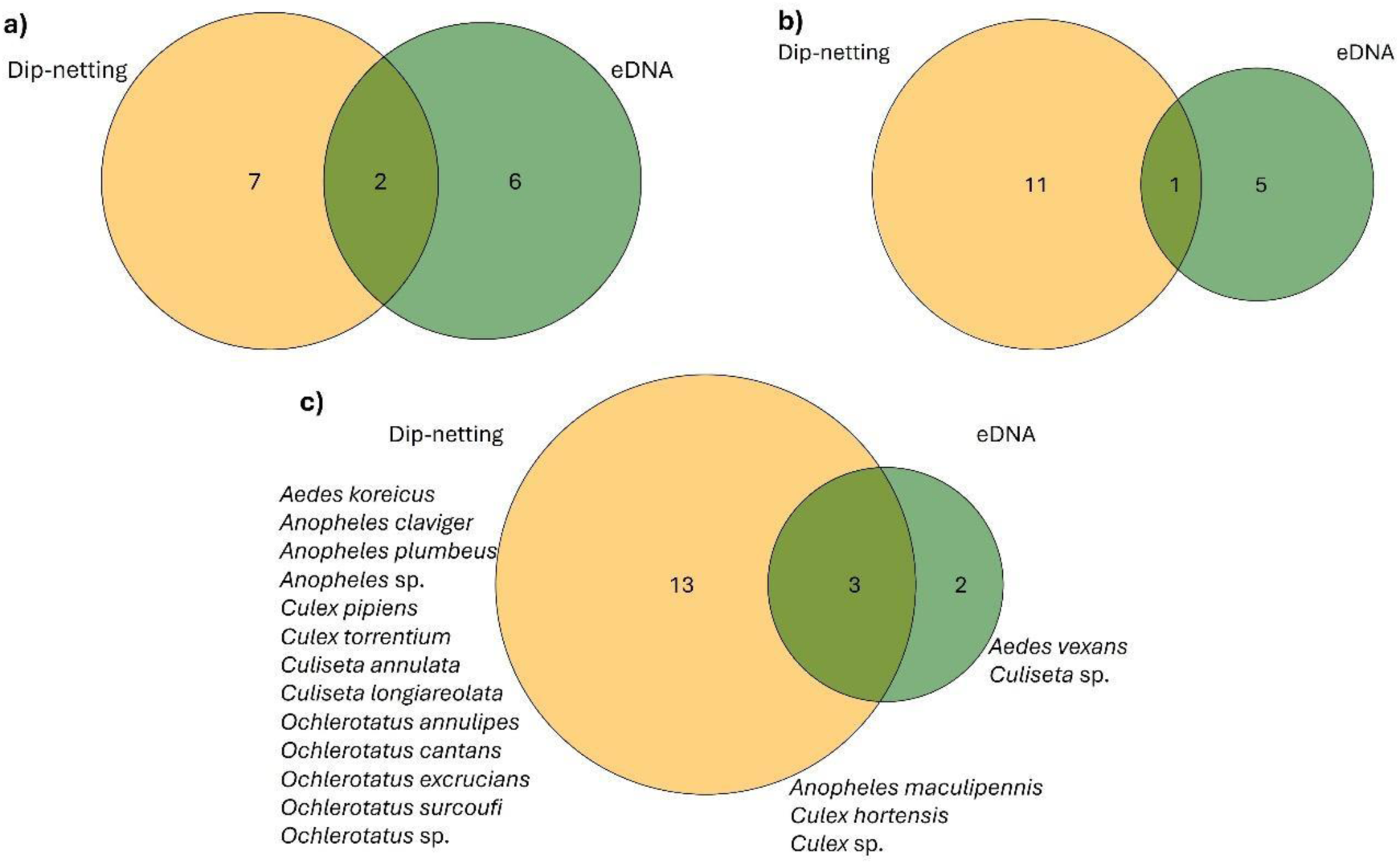
Venn diagrams showing the overlap in mosquito detection between sampling techniques (dip-netting and eDNA). Panels a and b show the number of ponds with mosquito species detected for garden (a) and urban ponds (b).

**Fig. 4.**
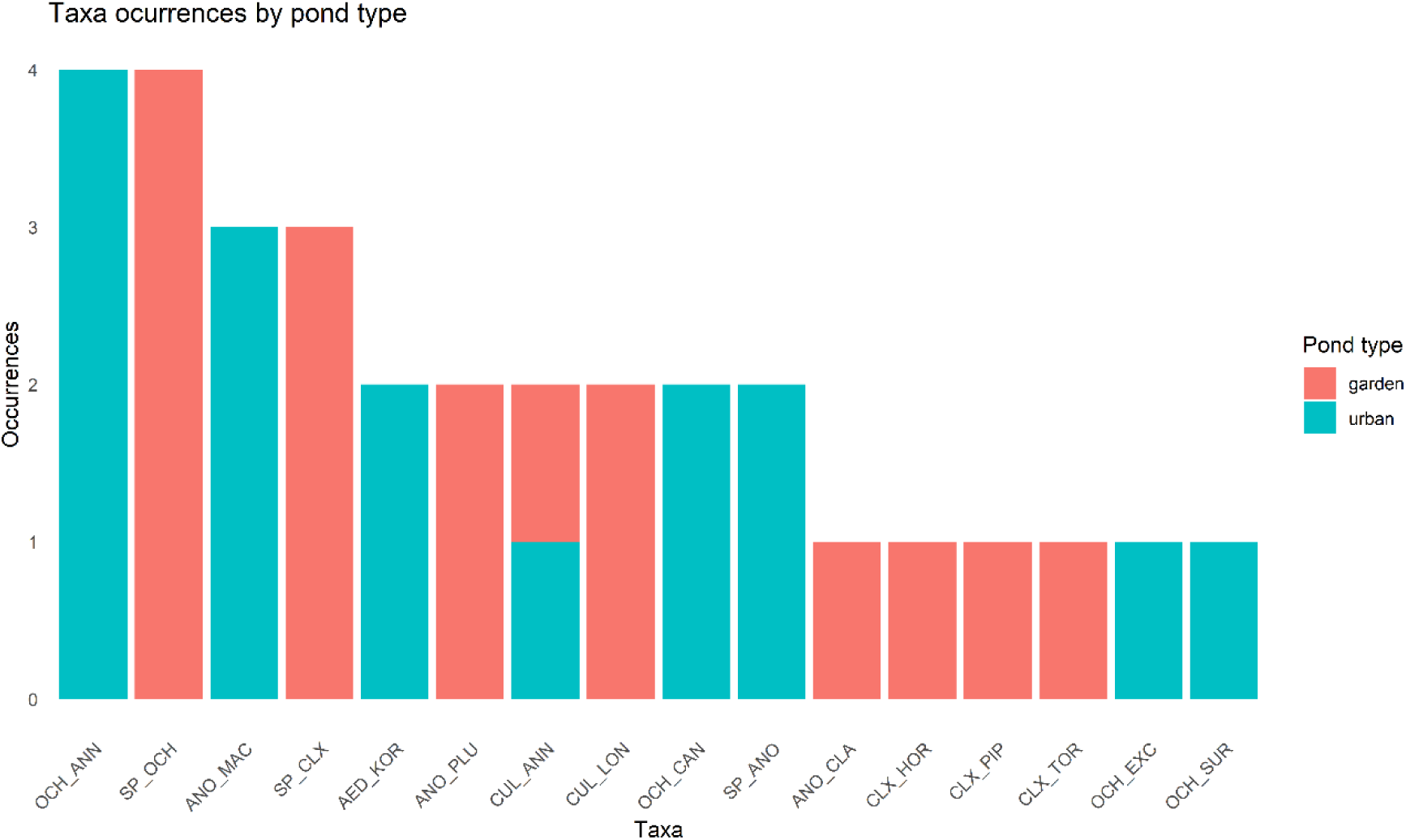
Barplot showing the mosquito taxa occurrence across pond types based on dip-net sampling. See Table 1 for taxa coding.

**Fig. 5.**
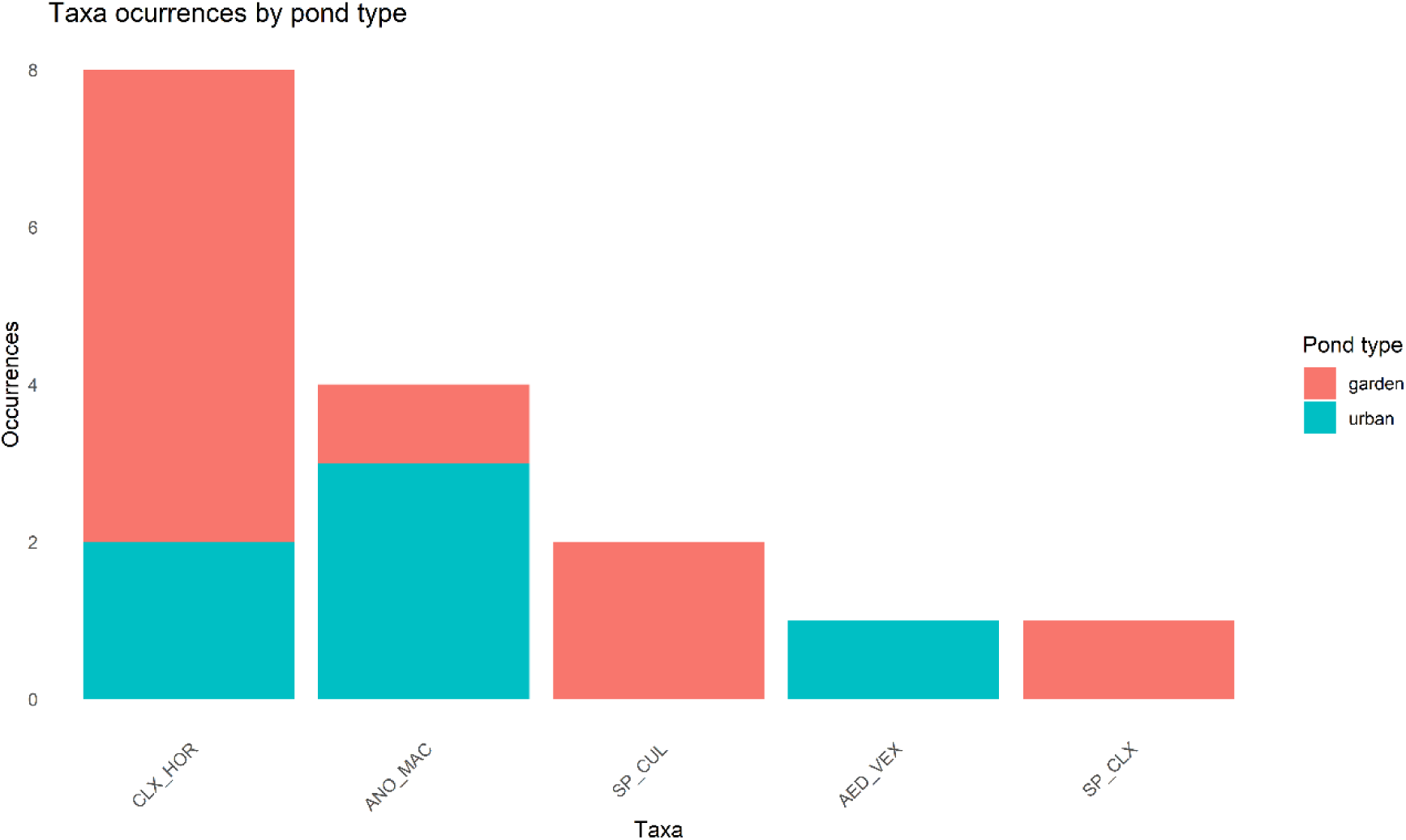
Barplot showing the mosquito taxa occurrence across pond types based on eDNA sampling technique. See Table 1 for taxa coding.

**Table 1.** Species of mosquito detected in garden and urban ponds based on our dip-netting sampling (left columns) and based on eDNA data (right columns). Numbers indicate the number of ponds in which each species was detected with each method. Species abbreviations used in figures are included.

|  | Dip-net data<br>from garden<br>ponds | Dip-net data<br>from urban<br>ponds | eDNA data<br>from garden<br>ponds | eDNA data<br>from urban<br>ponds |
| --- | --- | --- | --- | --- |
| *Aedes koreicus*<br>(AED_KOR) |  | 2 |  |  |
| *Aedes vexans*<br>(AED_VEX) |  |  |  | 1 |
| *Anopheles claviger*<br>(ANO_CLA) | 1 |  |  |  |
| *Anopheles maculipennis complex*<br>(ANO_MAC) |  | 3 | 1 | 3 |
| *Anopheles plumbeus*<br>(ANO_PLU) | 2 |  |  |  |
| *Culex hortensis*<br>(CLX_HOR) | 1 |  | 6 | 2 |
| *Culex pipiens*<br>(CLX_PIP) | 1 |  |  |  |
| *Culex torrentium*<br>(CLX_TOR) | 1 |  |  |  |
| *Culex* sp.<br>(SP_CLX) | 3 |  | 1 |  |
| *Culiseta annulata*<br>(CUL_ANN) | 1 | 1 |  |  |
| *Culiseta longiareolata*<br>(CUL_LON) | 2 |  |  |  |
| *Culiseta* sp.<br>(SP_CUL) |  |  | 2 |  |
| *Ochlerotatus annulipes*<br>(OCH_ANN) |  | 4 |  |  |
| *Ochlerotatus cantans*<br>(OCH_CAN) |  | 2 |  |  |
| *Ochlerotatus excrucians*<br>(OCH_EXC) |  | 2 |  |  |
| *Ochlerotatus surcoufi*<br>(OCH_SUR) |  | 1 |  |  |
| *Ochlerotatus* sp.<br>(SP_OCH) | 4 |  |  |  |

In the LM, fish presence had a significant negative effect, whereas MEM5 showed a marginally significant positive effect on mosquito abundance (Table 2), indicating that mosquito abundance was associated with spatial patterns at intermediate spatial scales (Figure S2).

**Table 2.** Result of a linear model using abiotic, biotic and spatial variables as predictors of variation in mosquito abundance.

|  | Estimate | SE | t | p |
| --- | --- | --- | --- | --- |
| (Intercept) | 0.122 | 0.071 | 1.731 | **0.087**· |
| Pond area | -0.014 | 0.020 | -0.714 | 0.477 |
| metalsPC1 | 0.013 | 0.022 | 0.619 | 0.538 |
| Fish presence | -0.117 | 0.042 | -2.816 | **0.006** ** |
| Amphibian presence | 0.058 | 0.041 | 1.408 | 0.163 |
| Predatory macroinvertebrates | -0.010 | 0.038 | -0.267 | 0.790 |
| Urbanisation | 0.016 | 0.103 | 0.159 | 0.874 |
| MEM2 | -0.040 | 0.028 | -1.426 | 0.157 |
| MEM5 | 0.037 | 0.019 | 1.950 | **0.054**· |

Statistical significance is indicated in bold as follows: · 0.1≤p; \*p≤0.05; \*\*p≤0.01

According to the GLM results explaining mosquito presence based on dip-net samples, both fish presence and MEM2 were negatively associated with the presence of mosquitoes, although MEM2 was only marginally significant (Table 3). MEM2 represents large-scale spatial patterns and distinguishes ponds located in the city centre from those in the suburbs (Figure S3). This pattern indicates a higher likelihood of mosquito presence in peripheral zones.

**Table 3.** Result of a generalised linear model using abiotic, biotic and spatial variables as predictors of variation in mosquito presence data based on dip-net sampling.

|  | Estimate | SE | z | p |
| --- | --- | --- | --- | --- |
| (Intercept) | 0.772 | 1.032 | 0.748 | 0.454 |
| metalsPC1 | 0.248 | 0.349 | 0.710 | 0.478 |
| Open water | -0.455 | 0.287 | -1.587 | 0.113 |
| Fish presence | -2.458 | 0.744 | -3.306 | **0.001** ** |
| Amphibian presence | -0.562 | 0.708 | -0.795 | 0.427 |
| Predatory macroinvertebrates | 0.092 | 0.535 | 0.171 | 0.864 |
| Urbanisation | -1.708 | 1.793 | -0.952 | 0.341 |
| MEM2 | -0.670 | 0.407 | -1.647 | **0.100**· |

Statistical significance is indicated in bold as follows: · 0.1≤p; \*p≤0.05; \*\*p≤0.01

In the GLM for mosquito eDNA-based presence, both urbanisation and MEM17 had a marginally significant positive effect on mosquito presence (Table 4). MEM17 reflects fine-scale spatial patterns (Figure S4).

**Table 4.** Result of a generalised linear model using abiotic, biotic and spatial variables as predictors of variation in mosquito presence data based on eDNA sampling.

|  | Estimate | SE | z | p |
| --- | --- | --- | --- | --- |
| (Intercept) | -2.977 | 1.028 | -2.897 | **0.004** ** |
| Open water | 0.667 | 0.531 | 1.257 | 0.209 |
| Fish presence | -1.117 | 0.683 | -1.634 | 0.102 |
| Amphibian presence | 0.382 | 0.712 | 0.537 | 0.591 |
| Predatory macroinvertebrates | 0.172 | 0.625 | 0.276 | 0.783 |
| Urbanisation | 2.647 | 1.456 | 1.819 | **0.069**· |
| MEM17 | 0.656 | 0.352 | 1.866 | **0.062**· |

Statistical significance is indicated in bold as follows: · 0.1≤p; \*p≤0.05; \*\*p≤0.01

Alternatively, we found a marginally significant positive interaction between fish presence and predatory macroinvertebrate abundance on mosquito abundance (Table S3, S4 and S5 in the Supplementary material).

According to the variance partitioning analysis, local abiotic (open water, metalsPC1, TN and pond area), local biotic (amphibians and fish presence, and abundance of predatory macroinvertebrates), and landscape (urbanisation and MEM2) fractions explained 7.8%, 4.9% and 3.6% of the total variance in mosquito community, respectively (Figure 5). Both the unique and shared fractions had significant effects on mosquito community composition.

**Fig. 5.**
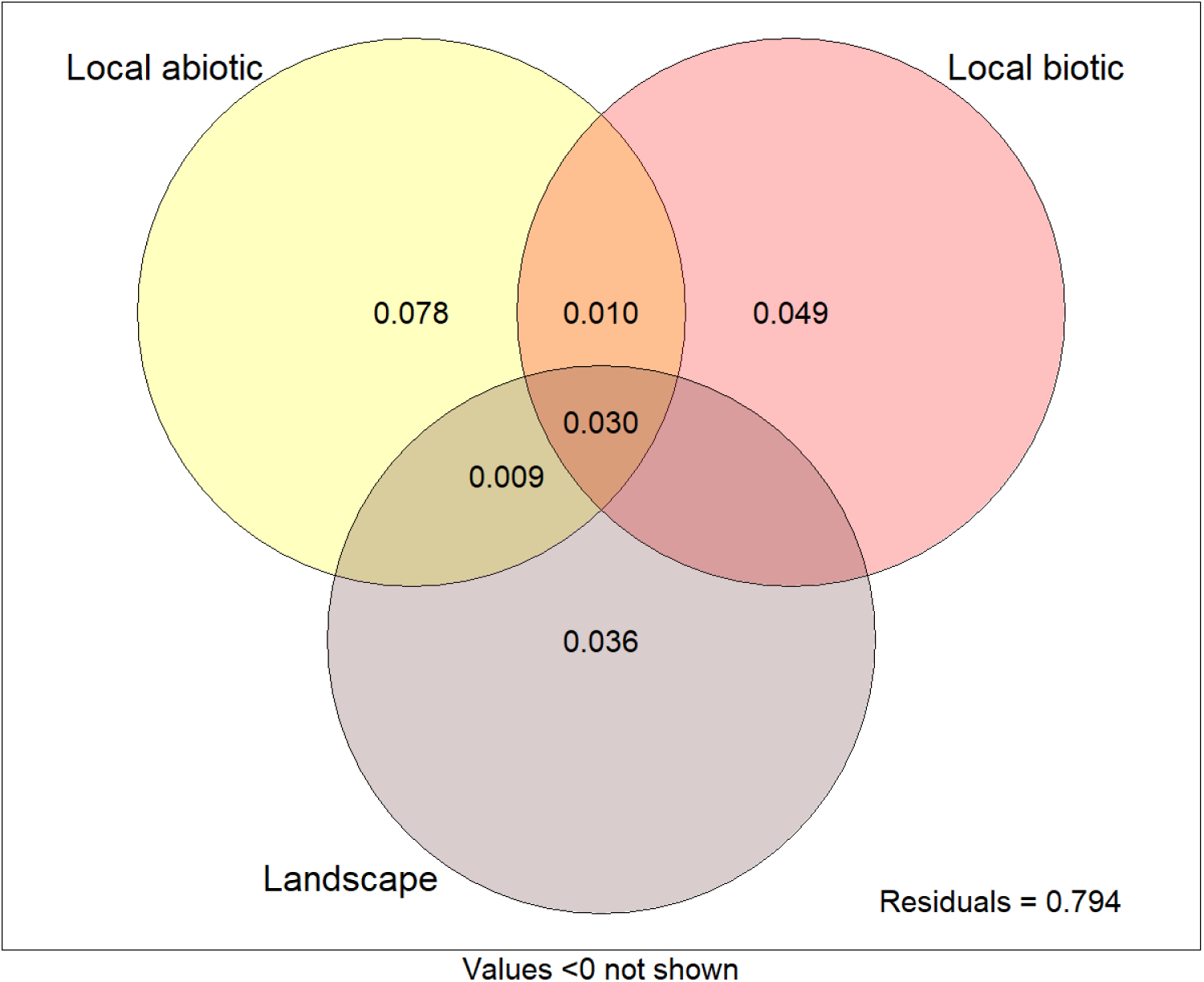
Venn diagram of the variation partitioning of the mosquito community explained by local abiotic, local biotic and landscape predictors.

## DISCUSSION

Here, we assessed whether urban ponds function as breeding habitats for mosquitoes in a Central European city, and examined the effects of potential drivers of mosquito occurrence and abundance. Overall, mosquitoes were detected in relatively few ponds - and generally in low abundance -, regardless of the type of pond or detection method. The presence of fish had the strongest negative effect on both mosquito abundance and presence, highlighting the importance of biotic interactions in shaping mosquito communities in urban ponds.

Despite using two complementary detection techniques, mosquitoes were detected only rarely. Individuals of the genus *Culex* were found mostly in garden ponds, with *Culex hortensis* being the only Culex species detected in urban ponds using eDNA. *Culiseta longiareolata* was recorded only in garden ponds, even though this species is typically associated with urban environments (Becker et al., 2010). In contrast, *Aedes* species were only found in urban ponds, together with most of the records of *Ochlerotatus*. These patterns suggest species-specific differences in pond type use. Increasing levels of urban pollution can reduce the quality of larval habitats, thereby promoting the dominance of genera such as *Culex*, known for their adaptability to temporary and degraded aquatic habitats (Carlson et al., 2004). In our study, *Culex hortensis* was the most frequently detected species, which has been previously shown to be positively influenced by landscape anthropisation (Perrin et al., 2023b). However, another study found this species mainly in natural habitats (Azari-Hamidian, 2007).

We found *Aedes koreicus* only in two of the sampled urban ponds. *Aedes vexans* was recorded in one urban pond, and was the only species exclusively detected by eDNA. Both *Aedes koreicus* and *Ae. vexans* are negatively influenced by landscape anthropisation (Perrin et al. 2023b; Perrin et al., 2022), which could explain the very low detection in our data. In urban settlements, *Ae. koreicus* generally shows habitat preference towards puddles, basin of fountains, or catch basins (Montarsi et al., 2013), habitats that were not included in our survey. In contrast, *Ae. vexans* prefers temporary floodwaters for breeding (Pothmann et al., 2025; Schäfer & Lundström, 2006), unlike other species of the same genus, such as *Ae. albopictus* or *Ae. aegypti* which prefer water-holding containers in urban environments (Herath et al., 2024; Kolimenakis et al., 2021).

We found *Anopheles plumbeus* in two garden ponds. This species is considered a potential malaria vector (Schaffner et al., 2012), with notes on its vector role for West Nile virus and dirofilariasis (Vermeil et al., 1960; Schaffner et al., 2001), but its low occurrence in our dataset suggests that these ponds are unlikely to represent important breeding habitats. A study performed in Belgium (Dekoninck et al. 2011), concluded that the strong population expansion of *An. plumbeus* was related to a larval habitat shift from tree-holes in natural ecosystems to anthropogenic influenced habitats, posing a public health concern with the expansion of this species into human-dominated environments. Another potential malaria vector is *An. claviger*, which we recorded in one of the garden ponds. This species has long been considered as a minor malaria vector due to its strong zoophilic tendency (Becker et al., 2010), but it has been increasingly detected in urban or peri-urban areas (Marí et al., 2014; Medlock & Vaux, 2014; Ruiz-Arrondo et al., 2023; Wegner, 2009) and has been associated with malaria outbreaks occurred in Eastern Mediterranean countries (Coluzzi et al., 1964; Gramiccia, 1956). The low detection of the species in our ponds could be due to its greater affinity for margins of ditches, canals, scrapes or small rocky puddles (Becker et al., 2010; Boukraa et al., 2016). The *An. maculipennis* complex was detected in urban ponds with both sampling methods and in garden ponds using eDNA. This species complex is typically associated with agricultural landscapes and is positively influenced by landscape anthropisation (Perrin et al., 2023b). High abundances of *An. maculipennis* complex have been reported from extensive breeding grounds, typically rice fields, and areas with farms but also urban areas (Gilioli et al., 2024). In other parts of Europe, this species has usually been collected in ephemeral and anthropogenic habitats such as ditches, irrigation channels and river bed pools (Calzolari et al., 2021; Kavran et al., 2018).

Consistent with Evans et al. (2019), we did not find a clear urbanisation effect on mosquito abundance. However, previous studies have reported differences in mosquito abundances across an urban gradient (e.g., Johnson et al., 2008; Li et al., 2014). Although spatial drivers in our study were only marginally significant, they do appear to suggest a trend that indicates a higher frequency of mosquitoes in ponds situated in suburban areas as opposed to those located in the city centre. Similar findings have been reported elsewhere, with Goselle et al. (2017) observing the highest abundance of several species in suburban areas. Furthermore, the meta-analysis of Perrin and collaborators (2022) found a general negative effect of urbanisation on mosquito abundance but also noted an increase in the abundance of mosquito species that are of global concern in human-modified landscapes. This suggests that urbanisation may favour a small subset of species which are adapted to anthropogenic landscapes.

Regarding the effect of heavy metals, although they were included as predictor variables in our analyses, we did not observe a significant effect on mosquito community composition, abundance or presence. The most abundant heavy metals in our ponds were manganese, aluminium, and iron, although it is important to remember that we studied their effect collectively, not individually. In agreement with our results, Mireji et al. (2008) did not find correlations between manganese or iron concentrations and mosquito species in urban aquatic habitats, while they found that copper was positively related to certain mosquito species. Notably, the average concentrations of these three heavy metals in our urban ponds were approximately two orders of magnitude lower than those in Mireji et al. (2008). Moreover, Ndenga et al. (2012) observed a negative relationship between *Anopheles* larvae abundance and iron content, but the iron concentrations in their study were much higher than those observed in our ponds. Regarding other water parameters, we did not observe a substantial effect of physico-chemical parameters on overall mosquito abundance, consistent with Beketov et al. (2010). Nevertheless, some species seem to have strong preferences for waters with high salinity and phosphate concentrations (Beketov et al., 2010), conductivity, dissolved oxygen, nutrients or pH (Avramov et al., 2024; Yadav et al., 2012). In addition, a recent meta-analysis by Avramov et al. (2024) has concluded that urban vector species are the most adaptable to a wider range of values for water quality properties.

Although variance partitioning revealed a relationship between the abundance of predatory macroinvertebrates and the mosquito community, we did not find a clear relationship between the abundance of mosquitoes and predatory macroinvertebrates in the regression analysis. Previous studies reporting a similar inconsistency have suggested that the presence of alternative prey may reduce predation on mosquito larvae, especially as the larvae grow and become harder to capture (Kumar et al., 2008). Complex submerged vegetation might also reduce overall predatory efficiency (Sunahara et al., 2002). However, we did not observe such a pattern. Possibly, in our ponds, macroinvertebrate predators’ abundance or diversity was not large enough to have a significant negative effect on the mosquito community, similar to findings from several studies that reported less diverse macroinvertebrate assemblages in artificial habitats, with a generally lower potential importance in limiting mosquito populations than natural habitats (Srivastava, 2006; Yanoviak, 2001).

In contrast to macroinvertebrate predators, fish presence in the ponds clearly reduced both mosquito abundance and presence, whereas amphibian presence had no such effect, even though they can also act as natural control agents (Perrin et al., 2023a). Fish introductions are common in urban ponds, often to control mosquitoes (Oertli & Parris, 2019). Fish may influence the numbers of mosquitoes both directly (through predation) and indirectly, by reducing the number of aquatic insects, which would otherwise be effective predators of mosquito larvae (Becker et al., 2010). Although some studies have reported a combined predatory effect of fish and macroinvertebrates on mosquito larvae (e.g. Hurlbert et al., 1972; Peck & Walton, 2008), in the present study, this effect was only marginally significant. However, the negative association between fish and mosquitoes should not be interpreted as evidence for fish stocking being an ecologically beneficial management solution, because fish can reduce amphibian populations, alter invertebrate assemblages, and shift pond ecosystems towards a turbid, phytoplankton-dominated state by reducing zooplankton biomass and the extent of macrophyte beds (Scheffer et al., 1993; Oertli & Parris, 2019; Bucciarelli et al., 2019; Hamer et al., 2002). Moreover, mosquito occurrence was relatively low across ponds overall, indicating that these habitats do not appear to function as major mosquito breeding sites in the studied urban landscape. Therefore, the ecological costs of fish introductions should be carefully considered in these systems.

Pond area did not have an effect on either mosquito abundance or presence in our study. This contrasts with previous empirical and experimental work reporting positive relationships between habitat size and mosquito abundance (Harlan & Paradise, 2006; Zhao et al. 2020). Other studies including both natural and artificial habitats (e.g. Westby & Juliano, 2017; Bradshaw & Holzapfel, 1988; Bevins, 2007; Sunahara et al., 2002) have found species-specific habitat size preferences. For instance, several studies reported that *Aedes japonicus* and *Culex* larvae preferred larger containers (Sunahara et al., 2002; Westby & Juliano, 2017), while *Ae. koreicus* preferred smaller habitats (Montarsi et al., 2013). While habitat size did not strongly influence total mosquito abundance in our ponds, it was one of the significant drivers of species composition, which is similar to findings by e.g. Talaga et al. (2017), where a significant fraction of the variation in community composition was explained by the size of the habitat, particularly at small spatial scales.

We observed differences between mosquito detection techniques related to pond type. In urban ponds, dip-netting performed better than eDNA, whereas in garden ponds, the detection rate was similar between techniques. Thus, dip-netting became overall much more efficient than eDNA since it allowed us to collect a larger number of taxa, and in a larger number of ponds. Moreover, eDNA surveys still need to be improved as the number of Culicidae species in the available databases is still somewhat limited, with only three genera (*Aedes*, *Anopheles* and *Culex*) accounting for 78% of the occurrences (Gutiérrez-López et al., 2023) and the databases are skewed in geographical and taxonomic average (Chapman et al., 2024). Despite these issues, it is a truly non-invasive method that does not harm species or habitats, and sometimes can have higher detection probabilities than traditional survey methods (Schneider et al., 2016). Given its comparable performance to dip-netting in garden ponds, eDNA could furthermore provide a useful tool for mosquito monitoring in citizen science programs. In our study, the lower eDNA detection rates in urban ponds may reflect dilution effects in the generally larger water bodies (Hansen et al., 2018; Tréguier et al., 2014).

While mosquito species composition was diverse across the ponds and the two pond types, their overall occurrence was relatively low, as mosquitoes were detected in only 32 of the 93 sampled ponds (34.4 %), and usually in low abundance. Nevertheless, some of the results should be interpreted with caution since the limited number of observations limit the robustness of the statistical associations. Furthermore, the early sampling season (spring), and the highly seasonal nature of mosquito assemblages in temperate regions might have contributed to the overall low occurrence and abundance of this group during our survey (Hanford et al., 2019; Rettich et al., 2007; Schäfer et al., 2004). While dip-net sampling captures only immature mosquitoes, which do not yet cause nuisance or spread diseases, their density can be used to predict the amount of adult mosquitoes (Zhao et al., 2020) and thus can provide certain relevant information that could be used in mosquito control at municipal level. In addition to ponds, other artificial water bodies, such as small plastic containers or flower pots in gardens, are also known to serve as mosquito breeding sites (European Centre for Disease Prevention and Control, 2012; Ibañez-Justicia, 2020), for which eDNA-based citizen science surveys could provide valuable data. Future comparative studies across different cities, climates, and mosquito habitats, and incorporating seasonality whenever possible, would provide a broader understanding of mosquito communities in urban ecosystems. Such knowledge is particularly important under ongoing climate change, as shifts in temperature, humidity, and precipitation can strongly influence mosquito distributions and the dynamics of mosquito-borne diseases (Brugueras et al., 2020; Boerlijst et al., 2023).

## Supporting information

Supplementary material_Tornero_preprint

## ACKNOWLEDGMENTS

We are grateful to all the citizen scientists, pond owners and managers who participated in the project. We acknowledge those who helped during the fieldwork and laboratory work: Zsuzsanna Márton, Beáta Szabó, Csaba F. Vad, Csilla Laskai, Péter Dobosy, Ádám Fierpasz, Kata Bene, Vivien Kardos, Bernadett Szabó, Virág Csiszár, Dóra Fehér, Tamás Felföldi, Bence Gergácz, Bence Buttyán, Khaulah Zakaria, Katalin Nagyné Bodolai, Károly Pálffy, Teresa Adhikary, and Varsha Rani.

