## Supplementary material_Tornero_preprint for "Drivers of mosquito presence and abundance in urban and garden ponds in a European city"

**Journal name:** Urban Ecosystems

**Author names:** Tornero, Irene; Barta, Barbara; Hamer, Andrew J.; Soltész, Zoltán; Huỳnh, Thu-Hương; Mészáros, Ádám; Horváth, Zsófia

Table S1. Garden ponds sweeps

| Garden ponds | |
|---|---|
| **Size range** | **Number of sweeps (aquarium net)** |
| Until 10 $m^2$ | 3 |
| 10 – 20 $m^2$ | 6 |
| 20 – 35 $m^2$ | 9 |
| > 35 $m^2$ | 10 with hand net |

Table S2. Urban ponds sweeps

| Other urban ponds | |
|---|---|
| **Size range** | **Number of sweeps (hand net)** |
| Until 500 $m^2$ | 10 |
| 500 – 4000 $m^2$ | 20 |
| 4000 – 40000 $m^2$ | 30 |
| > 40000 $m^2$ | 40 |

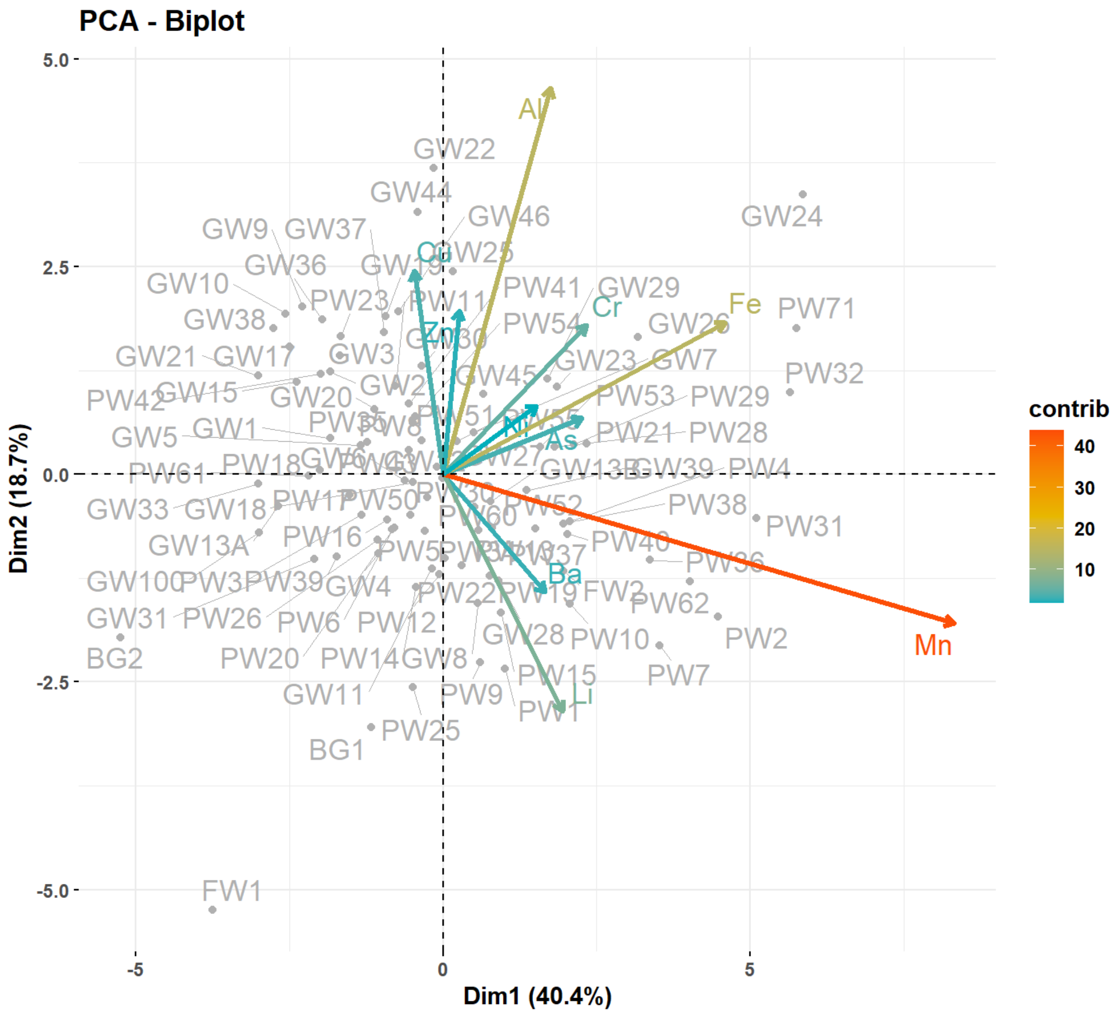


Figure S1. PCA biplot of the heavy metal concentrations in the 93 ponds.

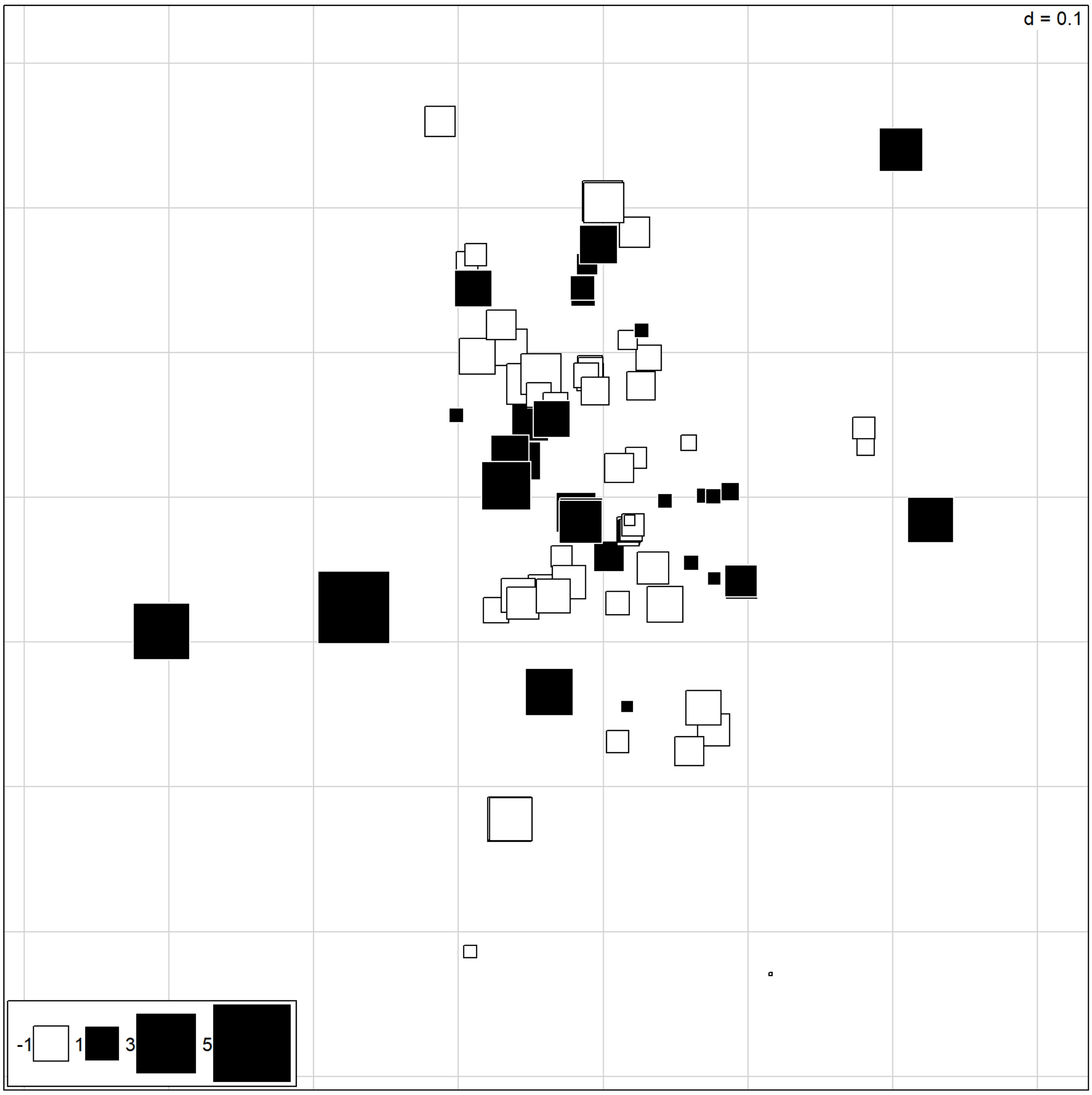


Figure S2. Moran's Eigenvector Maps, MEM5.

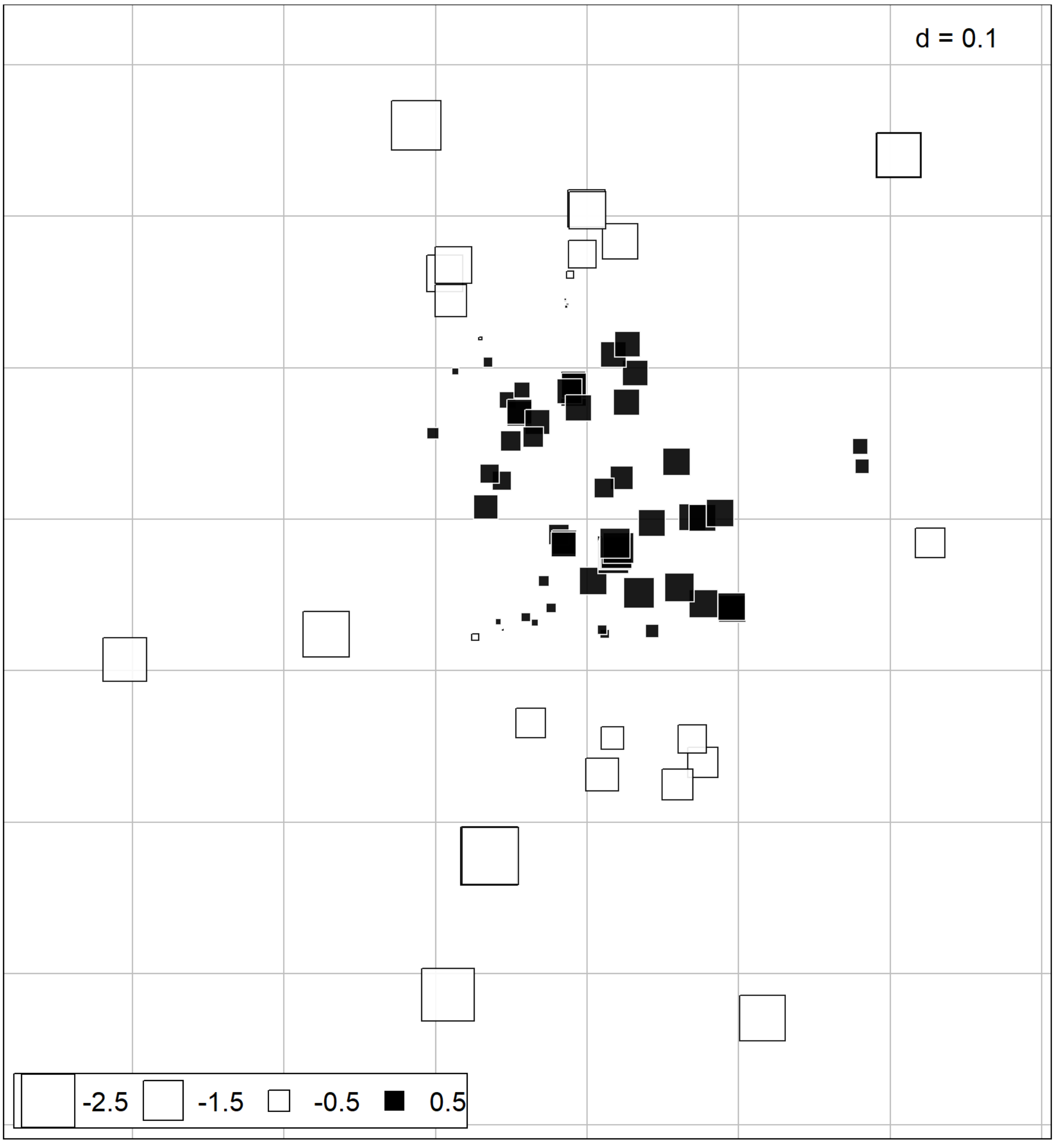


Figure S3. Moran's Eigenvector Maps, MEM2.

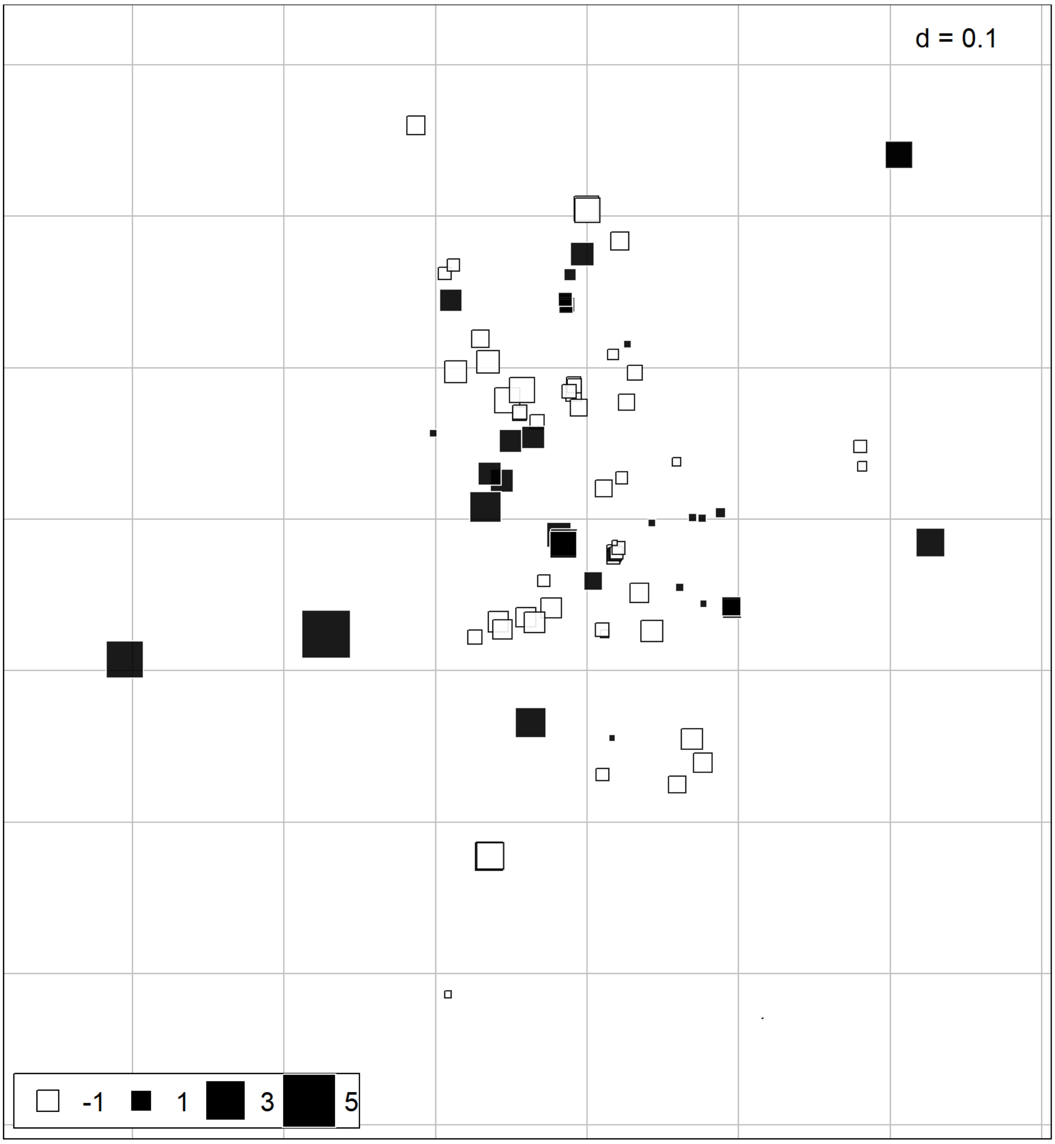


Figure S4. Moran's Eigenvector Maps, MEM17.

Table S3. Result of linear model using abiotic, biotic and spatial variables as predictors of variation in mosquito abundance.

| | Estimate | SE | t | p |
|---|---|---|---|---|
| (Intercept) | 0.167 | 0.075 | 2.237 | **0.028*** |
| Pond area | -0.015 | 0.020 | -0.767 | 0.446 |
| metalsPC1 | 0.006 | 0.022 | 0.259 | 0.796 |
| Fish presence | -0.161 | 0.049 | -3.312 | **0.001**** |
| Amphibian presence | 0.061 | 0.041 | 1.503 | 0.137 |
| Predatory macroinvertebrates | -0.233 | 0.137 | -1.703 | **0.092·** |

| | | | | |
|---|---|---|---|---|
| Urbanisation | 0.008 | 0.102 | 0.080 | 0.936 |
| MEM2 | -0.033 | 0.028 | -1.203 | 0.232 |
| MEM5 | 0.035 | 0.019 | 1.825 | **0.072·** |
| Fish presence * Predatory macroinvertebrates | 0.239 | 0.141 | 1.694 | **0.094·** |

Statistical significance is indicated in bold and as follows: · 0.1≤p; *p≤0.05; **p≤0.01

Table S4. Result of generalised linear model using abiotic, biotic and spatial variables as predictors of variation in mosquito presence data based on dip-net sampling.

| | Estimate | SE | z | p |
|---|---|---|---|---|
| (Intercept) | 0.779 | 1.146 | 0.680 | 0.497 |
| metalsPC1 | 0.246 | 0.370 | 0.665 | 0.506 |
| Open water | -0.456 | 0.287 | -1.586 | 0.113 |
| Fish presence | -2.463 | 0.831 | -2.965 | **0.003**** |
| Amphibian presence | -0.562 | 0.707 | -0.795 | 0.427 |
| Predatory macroinvertebrates | 0.069 | 1.942 | 0.035 | 0.972 |
| Urbanisation | -1.712 | 1.824 | -0.938 | 0.348 |
| MEM2 | -0.669 | 0.415 | -1.613 | 0.107 |
| Fish presence * Predatory macroinvertebrates | 0.025 | 1.996 | 0.012 | 0.990 |

Statistical significance is indicated in bold and as follows: · 0.1≤p; *p≤0.05; **p≤0.01

Table S5. Result of generalised linear model using abiotic, biotic and spatial variables as predictors of variation in mosquito presence data based on eDNA sampling.

| | Estimate | SE | z | p |
|---|---|---|---|---|
| (Intercept) | -3.072 | 1.045 | -2.940 | **0.003**** |
| Open water | 0.698 | 0.544 | 1.284 | 0.199 |
| Fish presence | -0.776 | 0.841 | -0.923 | 0.356 |
| Amphibian presence | 0.319 | 0.715 | 0.446 | 0.656 |
| Predatory macroinvertebrates | 1.622 | 2.168 | 0.748 | 0.454 |
| Urbanisation | 2.378 | 1.505 | 1.580 | 0.114 |

| | | | | |
|---|---|---|---|---|
| MEM17 | 0.561 | 0.376 | 1.489 | 0.136 |
| Fish presence * Predatory macroinvertebrates | -1.634 | 2.355 | -0.694 | 0.488 |

Statistical significance is indicated in bold and as follows: · 0.1≤p; *p≤0.05; **p≤0.01
